## Supplementary Information for "RNA secondary structure prediction using deep learning with thermodynamic integration"

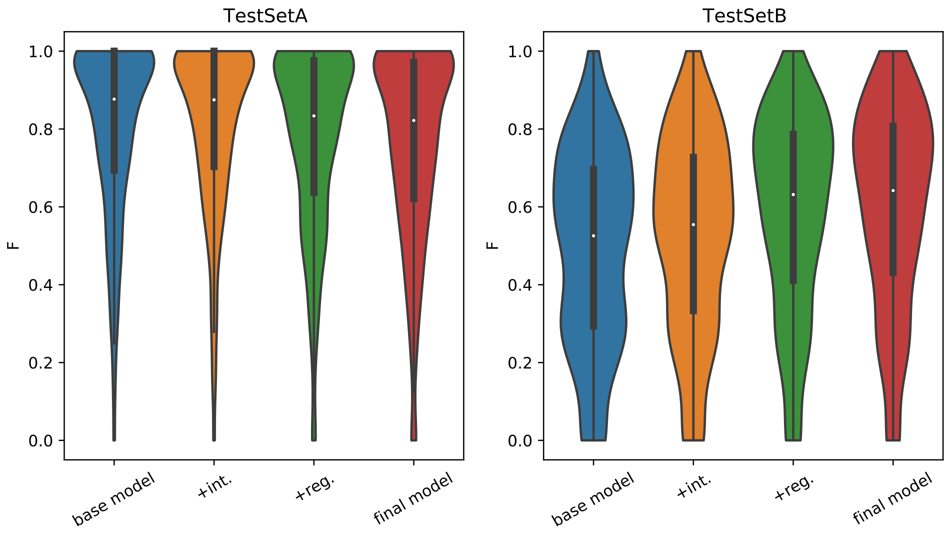


Supplementary Figure 1: $F$-values of the base model and the use of the thermodynamic-related techniques. +int.: the use of the thermodynamic-integrated folding scores, +reg.: the use of the thermodynamic regularization.

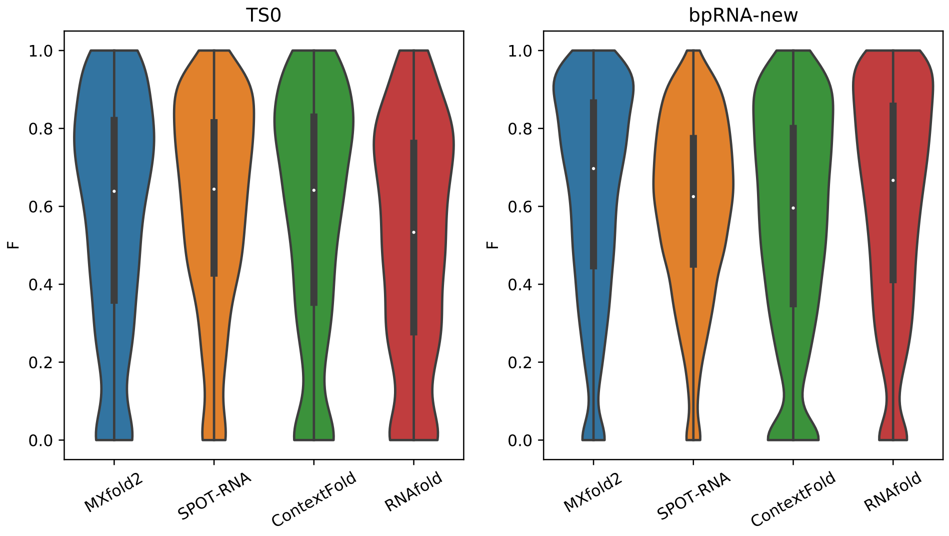


Supplementary Figure 2: $F$-values of MXfold2, SPOT-RNA, ContextFold, and RNAfold on the TS0 dataset for sequence-wise cross-validation (CV) and the bpRNA-new dataset for family-wise CV. All trainable methods were trained with the TR0 dataset.

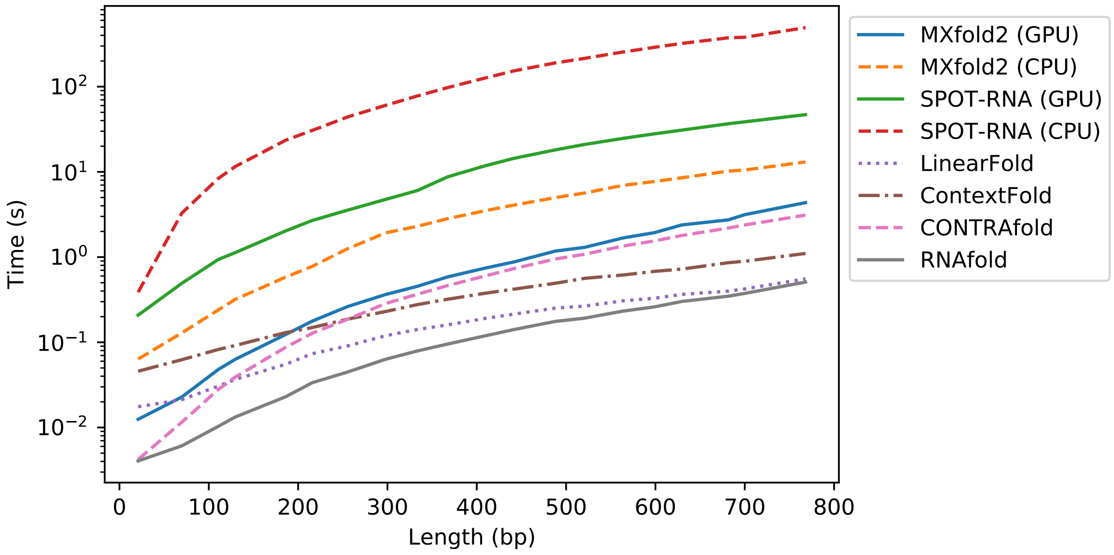


Supplementary Figure 3: The running time for the lengths of input sequences in TestSetA measured on Linux OS v4.15.0 with Intel XeonE5-2698v4 (2.20 GHz) and NVIDIA Tesla V100.

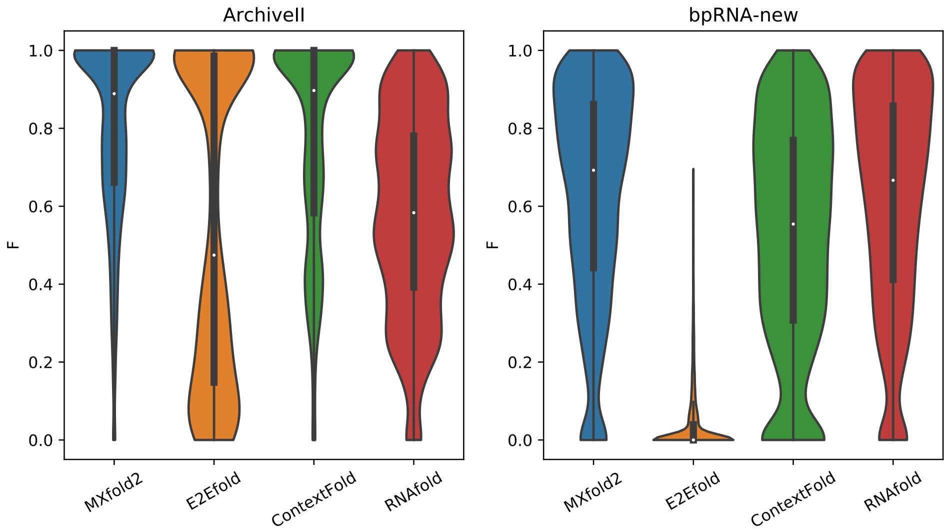


Supplementary Figure 4: $F$-values of MXfold2, E2Efold, ContextFold, and RNAfold using the ArchiveII dataset for sequence-wise cross-validation (CV) and the bpRNA-new dataset for family-wise CV. All trainable methods were trained using the RNAStrAlign dataset.

Supplementary Table 1: The summary of datasets used in our experiments.

| | | # sequences | length | source |
|---|---|---|---|---|
| Rivas Dataset | TrainSetA | 3,166 | 10–734 | Rivas *et al.* [3] |
| | TestSetA | 592 | 10–768 | |
| | TestSetB | 430 | 27–244 | |
| bpRNA-1m | TR0 | 10,814 | 33–498 | Danaee *et al.* [2] |
| | TS0 | 1,305 | 22–499 | |
| bpRNA-new | | 5,401 | 33–489 | the present study |
| RNAStrAlign | train | 20,923 | 30–600 | Tan *et al.* [5] |
| ArchiveII | | 3,966 | 28–1,800 | Sloma *et al.* [4] |
| T-Full | | 1,291 | 9–55 | Andronescu *et al.* [1] |

Supplementary Table 2: Comparison of the accuracy of the secondary structure prediction with the existing methods using TestSetA, TestSetB and the combined dataset with TestSetA and TestSetB.

| | TestSetA | | | TestSetB | | | Combined | | |
|---|---|---|---|---|---|---|---|---|---|
| | PPV | SEN | *F* | PPV | SEN | *F* | PPV | SEN | *F* |
| CONTRAfold | 0.708 | 0.745 | 0.719*** | 0.530 | 0.640 | 0.573*** | 0.633 | 0.701 | 0.658*** |
| CentroidFold | 0.701 | 0.678 | 0.678*** | 0.497 | 0.556 | 0.518*** | 0.615 | 0.627 | 0.611*** |
| ContextFold | 0.777 | 0.750 | **0.759** | 0.485 | 0.534 | 0.502*** | 0.654 | 0.659 | 0.651*** |
| LinearFold | 0.658 | 0.667 | 0.642*** | 0.501 | 0.609 | 0.544*** | 0.592 | 0.643 | 0.600*** |
| MXfold | 0.768 | 0.731 | 0.739*** | 0.561 | 0.620 | 0.582* | 0.681 | 0.684 | 0.673*** |
| MXfold2 | 0.754 | 0.778 | **0.761** | 0.571 | 0.650 | **0.601** | 0.677 | 0.724 | **0.693** |
| RNAfold | 0.658 | 0.668 | 0.642*** | 0.498 | 0.606 | 0.540*** | 0.590 | 0.642 | 0.599*** |
| RNAstructure | 0.657 | 0.650 | 0.631*** | 0.480 | 0.588 | 0.522*** | 0.582 | 0.624 | 0.585*** |
| SimFold | 0.654 | 0.643 | 0.629*** | 0.512 | 0.611 | 0.551*** | 0.594 | 0.629 | 0.596*** |
| Tornado | 0.749 | 0.754 | 0.746*** | 0.528 | 0.594 | 0.552*** | 0.656 | 0.687 | 0.664*** |

All trainable methods were trained using TrainSetA. For each test dataset, $F$-values that are significantly worse than the best are marked with * ($P < 0.05$), ** ($P < 0.01$) and *** ($P < 0.001$) as calculated with one-sided Wilcoxon signed-rank test. Others are in bold.

Supplementary Table 3: Comparison of the accuracy of the secondary structure prediction among MXfold2, E2Efold, ContextFold, and RNAfold.

| | Sequence-wise CV[1] | | | Family-wise CV[2] | | |
|---|---|---|---|---|---|---|
| | PPV | SEN | *F* | PPV | SEN | *F* |
| MXfold2[3] | 0.790 | 0.815 | 0.800 | 0.575 | 0.712 | 0.628 |
| E2Efold[3] | 0.605 | 0.519 | 0.548 | 0.0474 | 0.0307 | 0.0361 |
| ContextFold[3] | 0.770 | 0.771 | 0.768 | 0.559 | 0.506 | 0.522 |
| RNAfold | 0.551 | 0.613 | 0.577 | 0.552 | 0.720 | 0.617 |

[1] Sequence-wise cross-validation (CV) with the ArchiveII dataset.
[2] Family-wise CV with the bpRNA-new dataset.
[3] All trainable methods were trained using the RNAStrAlign dataset.
